# Plant-like heliotropism in a photosymbiotic animal

**DOI:** 10.1101/2023.11.02.565328

**Authors:** Eliska Lintnerova, Callum Shaw, Matthew Keys, Colin Brownlee, Vengamanaidu Modepalli

## Abstract

Being photosynthetic sessile organisms, plants established heliotropism to track the sun’s position across the sky and allow their vegetative parts to orient accordingly. Here, we report plant-like heliotropic movement in a photosymbiotic sea anemone *Anemonia viridis*. Like plants, photosynthesis represents a key energy source in endosymbiotic cnidarians bearing microalgae. We observed that *A. viridis* in their natural habitats under sunlight displayed heliotropism or solar tracking by pointing their tentacles towards the sun while remaining sessile, facing east at dawn and west at dusk as they track the sun’s relative position through the day, a phenomenon previously only observed in plants. Solar tracking movements in *A. viridis* are driven by a light wavelength that prompts photosynthesis in their endosymbionts. The heliotropic response was absent in both bleached (aposymbiotic) *A. viridis* and in symbiotic *A. viridis* with chemically inhibited photosynthesis. We revealed a direct correlation between heliotropism and endosymbiont oxygen production in *A. viridis*. Our findings suggest that photosymbiotic *A. viridis* has likely evolved plant-like heliotropism as an effective way to modulate exposure to solar irradiation for photosynthesis. The study exemplifies how photosynthetic organisms such as plants and photosymbiotic sea anemones, display similar behaviour in response to similar environmental pressures.

## Background

Sunlight is fundamental for most aspects of life on Earth, and it has been documented to induce physiological responses in various life forms. As for photosynthetic organisms like plants and algae, sunlight is an undeniably crucial factor for their survival [1]. As a result, plants and algae adapt and grow accordingly to optimise the amount of light reaching their surface for photosynthesis. For instance, motile green algae like *Chlamydomonas reinhardtii* can move towards light to increase photosynthesis **(Fig. 1A)** [2-4]. On the other hand, plants are sessile and restricted to the substrate where they germinated. Therefore, to enhance their photosynthesis, the plants evolved phototropism to re-orient the shoot growth towards a direction of the sunlight **(Fig. 1B)** [5-7]. Phototropism-like behaviour has been observed in other sessile organisms, including fungi and animals; for example, *Mucor circinelloides* and *Pilobolus crystallinus* fungi bending and growing their fruiting bodies towards blue light [8, 9]. Similarly, cnidarians such as polyps of *Eudendrium* (Hydrozoa), display plant-like phototropism under the blue light spectrum; the newly regenerated polyps of *Eudendrium* grew in the direction of the light source **(Fig. 1C)** and suggested that this process is comparable to bending of seedlings or a stem of plants **(Fig. 1B)** [10-12].

**Figure. 1:**
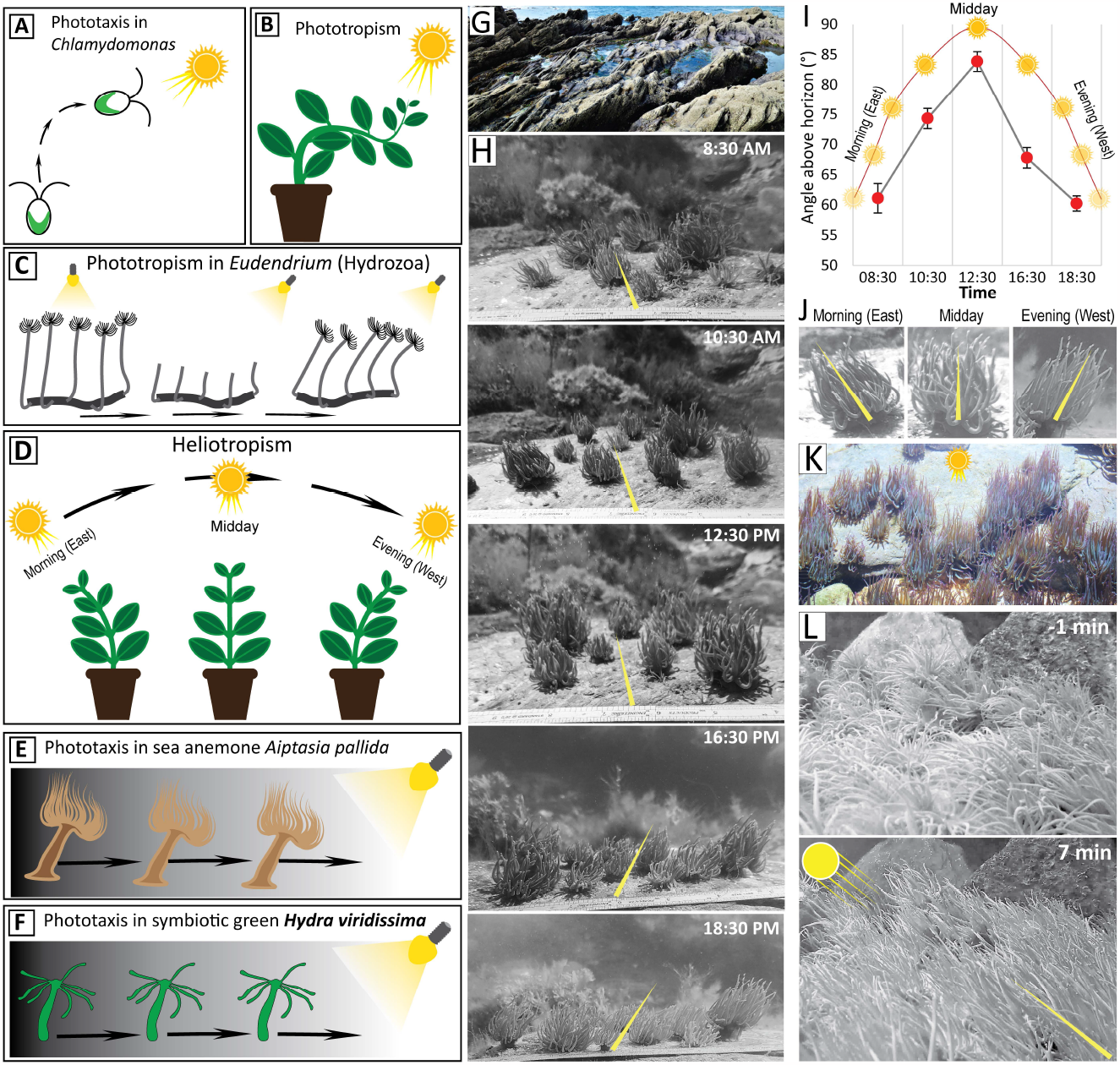
Illustration showing different light-mediated behaviours and evidence of heliotropism in *A. viridis*. **A**. Positive phototaxis behaviour in *Chlamydomonas reinhardtii*. **B**. Plants display phototropism, which is the growth in the direction of light. **C**. Newly regenerated polyps of *Eudendrium* tend to grow in the direction of the light source. **D**. Plants display heliotropism by tracking the sun’s path through the day. **E**. Positive phototaxis behaviour in *Aiptasia pallida* displayed by moving towards the light source. **F**. Positive phototaxis behaviour in *Hydra viridissima*. **G**. Rockpool habitat of *A. viridis*. **H**. Images of *A. viridis* collected across the day in the rockpool under sunlight. **I & J**. Change in the orientation of tentacles across the day, facing east at dawn and west at dusk, n≥7. **K**. *A. viridis* tentacles point towards the sun. **L**. Rapid response of *A. viridis* tentacles to restored sunlight.

Plants, in addition to phototropism like long-term growth responses to light, also display a more dynamic and oscillatory form of tropism known as heliotropism or solar tracking [13, 14], whereby plant leaves and/or flowers move to track the daily position of the sunlight through the day **(Fig. 1D)**. Heliotropic movement has been found in many plant species, regulating the amount solar irradiance absorbed for photosynthesis [15-17].

In some groups of animals, such as corals and sea anemones (Anthozoa, Cnidaria), phototactic responses are associated with a symbiotic relationship with photosynthetic algae. Dinoflagellates of the genus *Symbiodinium* (also known as zooxanthellae) reside as endosymbionts in the cnidarian host cells [18-21]. Similar to plants, photosynthesis represents an important energy source in endosymbiotic cnidarians [22], and thus, the amount of solar irradiation reaching organisms is essential and possibly drives complex light-driven responses similar to plants. Symbiotic corals were shown to develop different morphology based on the irradiance available in their habitat; they can grow flattened in the low light environment or more branched in the environment with higher irradiance to create self-shading. Additionally, the coral tissue and skeletal plates can become thinner in the environment with lower irradiance [23-25]. Likewise, sea anemones exhibit several distinct behavioural responses towards the light. Anemones can expand or contract the oral disk and flex or retract the tentacles depending on the light conditions. Though anemones are mainly sedentary, they do move across the surface. Anemones like *Aiptasia pallida* **(Fig. 1E)** display phototaxis behaviour by moving towards or away from a light source [18, 26, 27]. Similarly, in Hydra (Cnidaria, Hydrozoan), symbiotic green *Hydra viridissima* **(Fig. 1F)** and asymbiotic brown *Hydra vulgaris* display phototaxis [18, 28-31]. Nevertheless, it is unclear how these light-dependent responses are manifested in behavioural adaptations to natural habitats and the lifestyle of photosynthetic anemones, and we lack an in-depth understanding of their behaviour under sunlight.

From laboratory experiments, It is clear that cnidarians exhibit vivid responses to light, either by growing towards the light (phototropism) **(Fig. 1C)** or by moving towards or away from a light source (phototaxis) **(Fig. 1E & F)** [18, 28-30]. Being sessile, plants established heliotropism or solar tracking **(Fig. 1D)**, a strategy to track the sun’s position throughout the day, a phenomenon unique to plants [13, 14]. Here, we aim to determine definitively whether photosymbiotic anemones exhibit plant-like heliotropism, which means whether they can track the sun’s position throughout the day by pointing their tentacles towards the sun while remaining sessile. If this is the case, can they respond to changes in sunlight intensity and modulate their behaviour accordingly? Furthermore, we aim to test whether these changes in behaviour are directly associated with the photosynthesis of their endosymbionts through a series of experiments.

## Results and Discussion

### *A. viridis* exhibits heliotropism

Here, we examined the snakelocks sea anemone (*Anemonia viridis*), a photosymbiotic animal that belongs to the phylum Cnidaria and class Anthozoa. *A. viridis* occupies a wide range of light-exposed habitats and are largely dispersed in shallow rock pools with the highest mid-day solar irradiance **(Fig. 1G)** [32, 33]. Observing *A. viridis* in their natural habitats under sunlight revealed plant-like heliotropism by pointing their tentacles towards the sun, facing east at dawn and west at dusk as they track the sun’s relative position **(Fig. 1H-J)**. Temporary blocking of sunlight resulted in rapid loss of heliotropic behaviour by distributing the tentacles randomly **(Fig. 1K & L)** (**Movie S1)**.

We replicated the heliotropic response under laboratory conditions to gain insights into the role of photosynthesis in heliotropic behaviour. First, we analysed their behavioural responses to light and the impact of light quality on the tentacle orientation. Under laboratory conditions, within 5-7 minutes, individuals displayed maximum orientation towards light from above, with tentacles reaching the three highest sections, VIII, IX and X **(Fig. 2A & B)**. In the wild, *A. viridis* exhibits a strong heliotropic response on sunny days and the magnitude of the response corresponds to the strength of the sun radiation **(Fig. 1K)**. Like the field observations, the response magnitude was proportional to the intensity of light applied **(Fig. 2C & D, Movie S2)**. Next, we tested if *A. viridis* displayed a diaheliotropic response under laboratory conditions. Similar to natural sunlight tracking **(Fig. 1H-J)**, under laboratory conditions, *A. viridis* could track the movement of a unidirectional light source that was rotated by 90° at 10-minute intervals **(Fig. 1H, Movie S3)**. Thus, heliotropism in *A. viridis* is probably triggered by immediate light stimuli, as the tentacle’s orientation can adjust rapidly to changes in the light position.

**Figure. 2:**
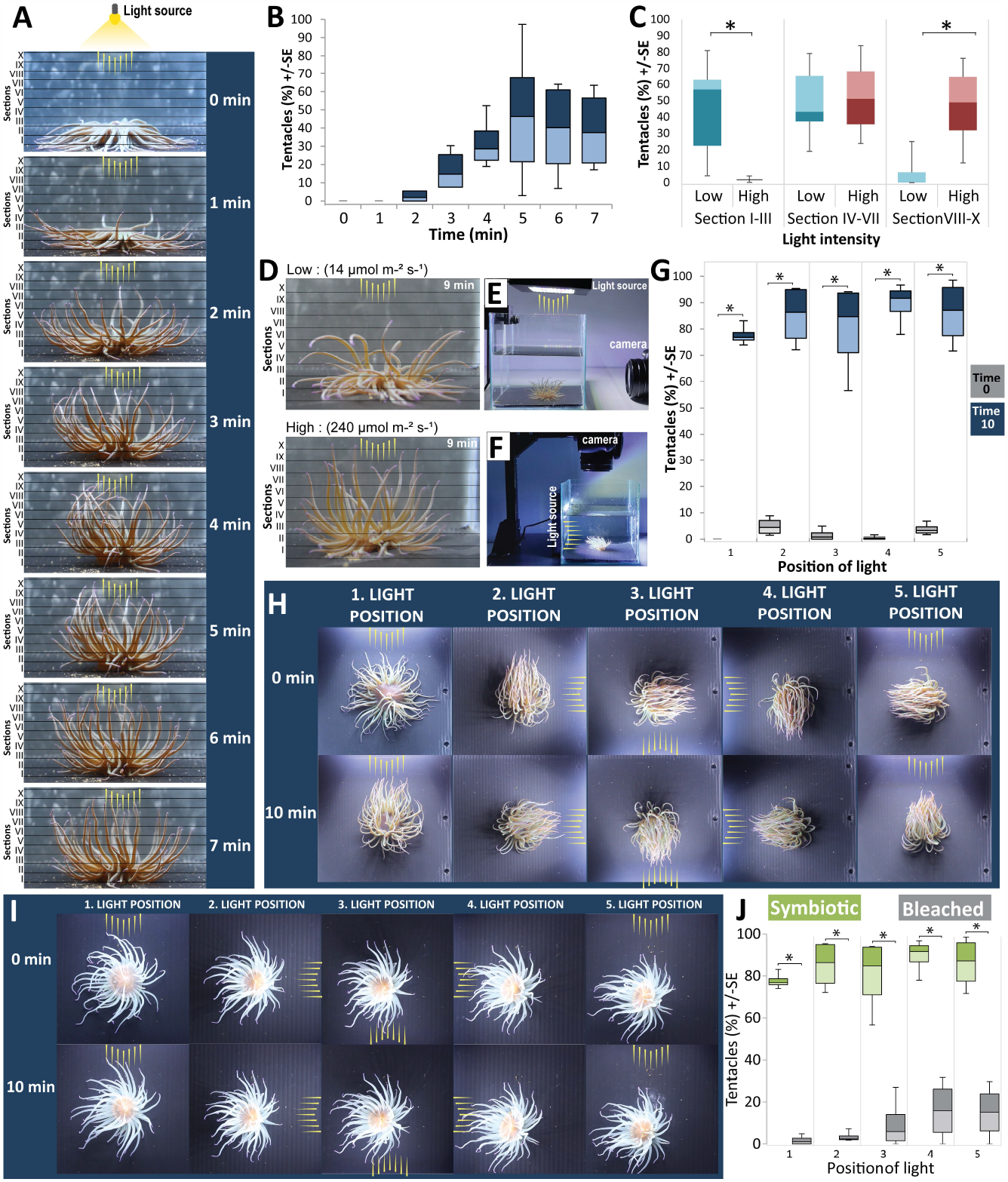
Heliotropism in *A. viridis* and its association with the photosynthetic endosymbionts. **A**. The response time of *A. viridis* tentacles to white light exposure from above (arrows) and the scale used to establish the position of tentacles (horizontal bars). **B**. The proportion of tentacles reaching the three highest sections, VIII, IX and X, over seven minutes of light exposure (Friedman test, X2(7) = 22.417, *p < 0.05, n=4). **C & D**. The *A. viridis* exhibits a strong response under bright light conditions (Section I-III; Section VIII-X, Paired T-test, *p < 0.05, n=6). **E**. The experimental setup used in A & D. **F**. The experimental setup used in H & I. **G**. The proportion of *A. viridis* tentacles pointing towards the light source positioned on different sides of the aquarium at 0 and 10 minutes of light exposure (For all the light positions 1-5, Paired T-test, *p < 0.05, n=4). **H**. Images of an *A. viridis* at the beginning and the end of five 10 minutes-long exposures to light positioned on the different sides of the aquarium. **I**. Images of an aposymbiotic (bleached) *A. viridis* at the beginning and the end of five 10 minutes-long exposures to light positioned on different sides of the aquarium. **J**. The proportion of symbiotic vs bleached *A. viridis* tentacles pointing towards the light source at the end of five 10 minutes-long exposures to light positioned on different sides of the aquarium (For all the light positions 1-5, Two sample T-test, *p < 0.05, n=4). The yellow arrow in panels A, D, E, F, H & I symbolises the direction of the light.

### Symbiotic *A. viridis* displays altered responses to different wavelengths of light

In plants, heliotropic responses are specific to violet and blue light (380-490 nm) [6, 7, 13, 14]. Likewise, we tested the response of *A. viridis* to different light spectra by comparing the percentage of tentacles reaching the two highest sections, IX and X, after 9 minutes of light exposure **(Fig. 2A)**. Both violet (380-435 nm) and blue (435-490 nm) wavelengths elicited the heliotropic response in *A. viridis*. In contrast, wavelengths in the cyan-green region (490-520 nm) were ineffective **(Fig. 1A & B, Movie S4)**. It is evolutionary important for the plant to sense and react to the blue light spectrum as it positively influences photosynthesis [34, 35]. Hence, symbiotic cnidarians gain a substantial part of their energy from photosynthetic *Symbiodinium* [22, 36], it is probably adapted to react to blue light as well. On the contrary, exposure to cyan had an insignificant effect on *A. viridis* heliotropic response. Cyan light consists mainly of a green spectrum, having the lowest impact on the photosynthetic rate [37, 38] and even on the phototactic responses in other symbiotic anthozoans [39, 40]. Therefore, similarly to vascular plants, triggering heliotropic behaviour in *A. viridis* by blue light suggests that photosynthesis (executed by *Symbiodinium*) is the key regulatory factor.

### Heliotropism in *A. viridis* allies with the presence of the endosymbiont

To determine the potential role of endosymbionts in the heliotropic response, we tested the heliotropic response of bleached (aposymbiotic) *A. viridis* **(Fig. 3E-H)**. Compared to symbiotic *A. viridis* **(Fig. 3A & B)**, bleached *A. viridis* **(Fig. 3C & D)**, devoid of photosymbionts **(Fig. 3E & H)**, showed a substantially reduced response to blue and violet light **(Fig. 3C & D, Movie S5)**. As for the symbiotic *A. viridis*, on average, 61.69% and 47.37% of tentacles pointed straight up after 9 minutes of blue and purple light exposure **(Fig. 3A)**. In contrast, only 4.51% of tentacles of bleached *A. viridis* rose to this level under blue exposure, and none under purple light exposure **(Fig. 3C & D, Movie. S4)**. Moreover, bleached A. *viridis* lost the ability to track the position of the light source **(Fig. 1I & J, Movie S6)**. Therefore, *Symbiodinium* plays an important role in the heliotropic response of *A. viridis*. These observations also coincide with the phototaxis response in *Aiptasia pallida* [18, 41] and the limited ability of bleached *A. viridis* to respond to light [19, 20, 26]. Therefore, the host responds to *Symbiodinium* mediated stimuli [18-20, 41], potentially photosynthetic-derived products, such as oxygen or organic compounds, signalling the behavioural response.

**Figure. 3:**
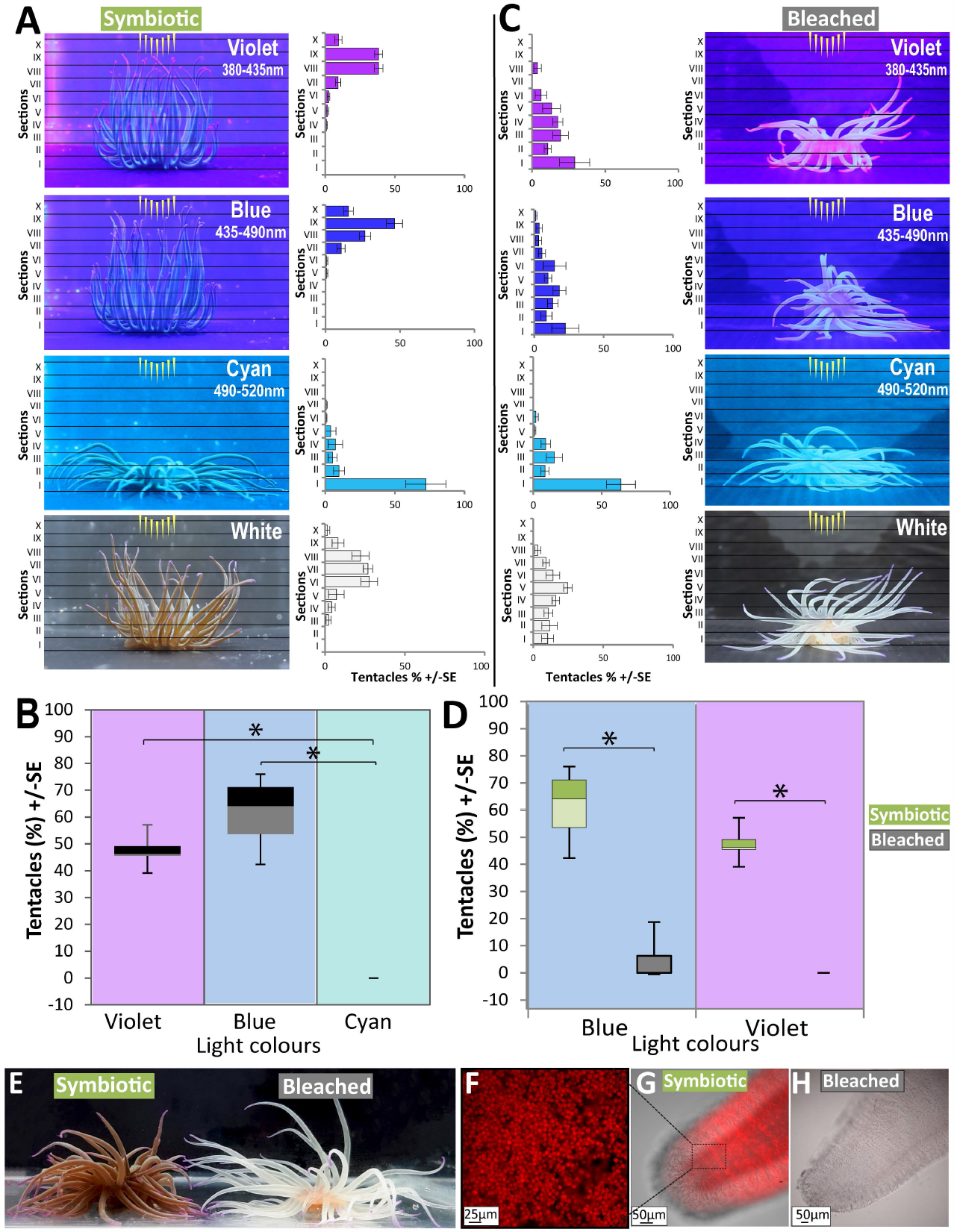
The effect of light wavelength on heliotropic response in symbiotic and bleached (aposymbiotic) *A. viridis*. **A**. The effect of light wavelength on symbiotic *A. viridis*. **B**. The proportion of symbiotic *A. viridis* reaching the two highest sections IX, X after 9 minutes of exposure to different colours of light (One-way repeated measures ANOVA, F_4,20_=83.013, *p < 0.05. eta2[g]=0.898, n=6). **C**. The effect of light wavelength on bleached *A. viridis*. **D**. The proportion of symbiotic and bleached *A. viridis* reaching the two highest sections IX, X in response to light after 9 minutes of exposure to blue (435-490 nm) and violet (380-435 nm) treatment (Blue and purple light exposure, Wilcoxon rank sum test, W=0, *p < 0.05, n=6). **E**. Symbiotic and bleached *A. viridis*. **F**. *Symbiodinium* chlorophyll fluorescence inside a tentacle. **G & H**. Tentacle of symbiotic and bleached *A. viridis*, the *Symbiodinium* is absent in bleached *A. viridis*. The yellow arrows in panels A and C show the direction of the light source.

Interestingly, in bleached *A. viridis* the response to light does not completely disappear; instead, an unsystematic and low level of response was apparent irrespective of the light wavelengths (**Fig. 3C, Movie S5)**. The low level of response in bleached *A. viridis* implicates the involvement of photoreceptors such as opsin and cryptochrome, similar to the other light-responsive behaviours in cnidarians [42-44]. However, qPCR analysis of opsin and cryptochrome gene expression did not show any significant differences between symbiotic and bleached *A. viridis* **(Fig. 4A)**, indicating the presence of *Symbiodinium* is the crucial factor underlying the heliotropic response. The mild response of bleached *A. viridis* towards the light (**Movie S5)** could be solely mediated through *A. viridis* photoreceptors, which is similar to other asymbiotic or non-symbiotic anthozoan species that sense and respond to light [44, 45]. It could be reasoned that bleached *A. viridis* can sense the light; however, the lack of *Symbiodinium* specifically hinders the heliotropic response. Thus, it seems that there are two levels in symbiotic *A. viridis* sensing and responding to the incoming light: through the *A. viridis* itself and a secondary stimulus from its endosymbionts. Since bleached *A. viridis* tentacles show random movements (**Fig. 2C, Movie S5)**, it is likely the supply of photosynthesis-derived stimuli from endosymbiont underlies the directional heliotropic tentacle movement.

**Figure. 4:**
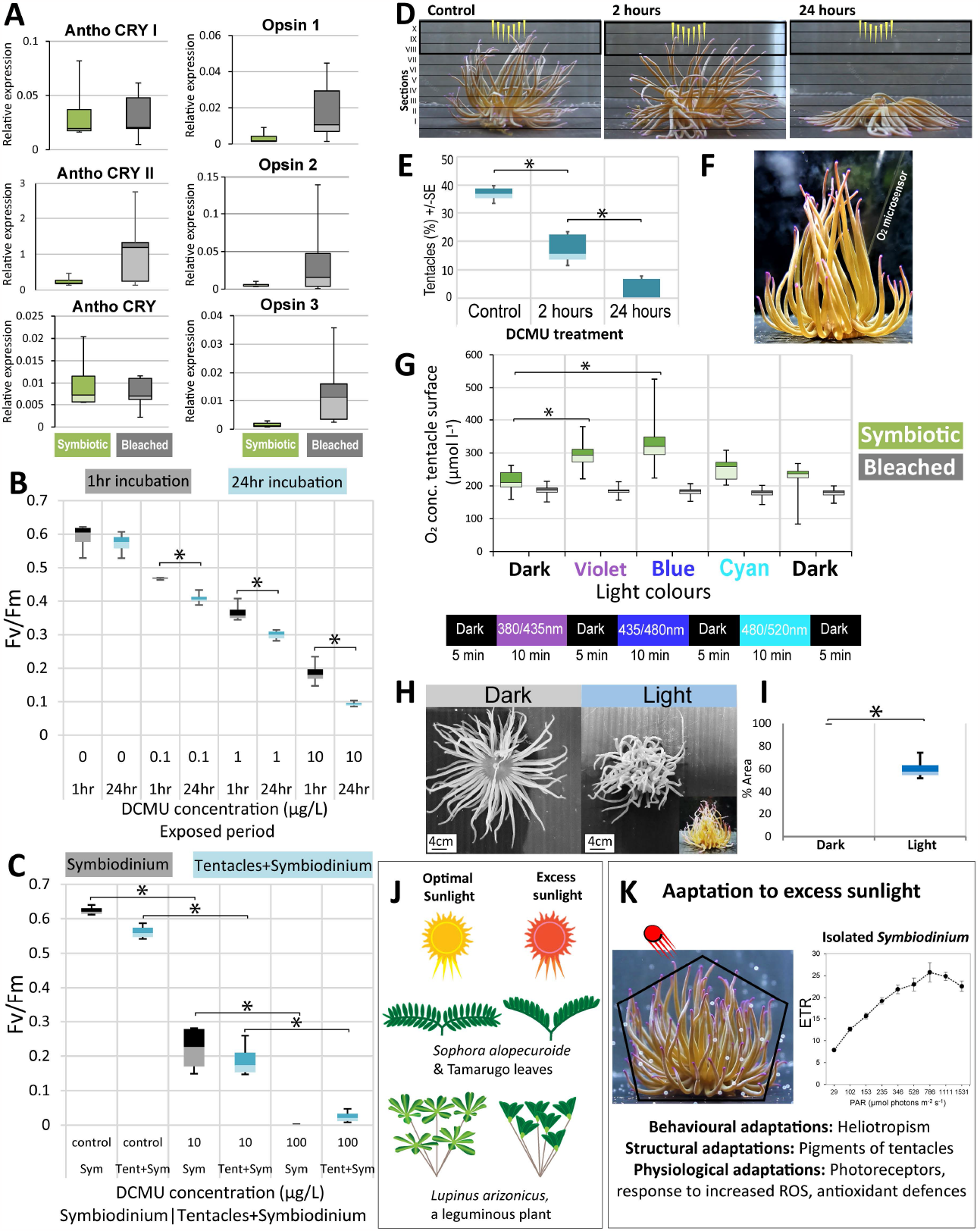
Heliotropism in *A. viridis* is directly associated with the photosynthesis of endosymbionts. **A**. Expression levels (Cq value assessed by qPCR) of candidate cryptochromes and opsin genes measured in tentacles of symbiotic and bleached *A. viridis* (n=5). **B**. Effect of different DCMU concentrations on Fv/Fm of *Symbiodinium* after 1 hour- and 24 hours-long exposure (Paired samples t-test, *p < 0.05). **C**. Effect of different DCMU concentrations on Fv/Fm of *Symbiodinium* and symbiotic tentacles after 1-hour exposure (Paired samples t-test, *p < 0.05). **D & E**. The proportion of *A. viridis* tentacles reaching the top sections VIII, IX and X after 5 minutes of light exposure either before (Control) or 2 hrs and 24 hrs after the addition of DCMU (One-way repeated ANOVA, F_2,6_ = 6.028, *p < 0.05, eta2[g]= 0.423). **F & G**. O_2_ measurements were collected at the surface of symbiotic and bleached *A. viridis* tentacles under different light conditions (Two-tailed test, *p < 0.05, n=4). **H & I**. Reduction of tentacle percentage area exposed to light during heliotropism (T-test, p < 0.05, n=4). **J**. Illustration showing paraheliotropism in plant species: *Lupinus succulentus* [50, 51], *Prosopis tamarugo* [52], *Sophora alopecuroides* [53]. **K**. Various adaptations of *A. viridis* to modulate influx solar irradiation. Instantaneous light curves of electron transfer rate (ETR) of *Symbiodinium* isolated from tentacles. Error bars indicate standard deviation (n ≥ 6).

### Heliotropic response in *A. viridis* is activated through photosynthesis of their endosymbionts

Our findings raise the key question of whether photosynthesis drives heliotropic responses in *A. viridis*. Therefore, we tested the correlation between endosymbiont photosynthesis with *A. viridis* heliotropic behaviour by inhibiting photosynthesis with the electron transport uncoupler 3-(3,4-diclorophenyl)-1,1-dimethyl urea (DCMU) [46]. DCMU is commonly used in generating aposymbiotic cnidarian hosts [46, 47]. A toxicology study on the effects of biocides on symbiotic and aposymbiotic juveniles of the coral *Acropora tenuis* revealed that exposure to DCMU in concentrations of 0.1, 1, 10 and 100 μg/L for ten days did not cause abnormalities such as tissue detachment or death in aposymbiotic juveniles, which suggests that DCMU is non-toxic to the anthozoan host [48]. The effects of DCMU on *Symbiodinium* have been extensively studied and have shown that their photosynthetic activity can be significantly affected at very low concentrations [47, 49]. Similarly, in our experiments, the DCMU treatment significantly reduced the mean photosynthetic efficiency (Fv/Fm) in both isolated *Symbiodinium* and *Symbiodinium* within the *A. viridis* tentacles in a time- and dose-dependent manner **(Fig 4B & C)**. The length of DCMU exposure had a negative effect on *A. viridis* ability to respond towards the light source, thus on their heliotropic behaviour and was completely lost in *A. viridis* exposed to DCMU for 24 hours **(Fig 4D & E)**. Likewise, the use of DCMU negatively influences the phototactic behaviour of *Anthopleura elegentassima* (in this case, expansion and retraction of the tentacles) [19]. Together, these results suggest that DCMU exposure inhibited the heliotropic response of *A. viridis* **(Fig 4D & E)**, emphasising the central role of photosynthesis in the *A. viridis* heliotropic response.

Next, to test the association between *A. viridis* heliotropism and the rate of photosynthesis, we measured O_2_ production by *Symbiodinium*. O_2_ concentrations on the surface of *A. viridis* tentacles were monitored with an O_2_ microsensor at different light conditions **(Fig 4F & G)**. Exposure to violet and blue wavelengths (provoking the strong heliotropic response) resulted in a significant O_2_ production compared to cyan wavelengths and no-light conditions (that do not drive the heliotropic response). This further highlights the direct correlation between the rate of photosynthesis and the *A. viridis* heliotropic response. As expected, bleached *A. viridis* showed insignificant O_2_ production under any light conditions. In the absence of light, there was no significant difference between symbiotic and bleached *A. viridis* **(Fig 4G)**.

### Is heliotropism in *A. viridis* an adaptive mechanism to promote light absorption or conversely to minimise excess light uptake?

Together, our experiments show that photosynthesis drives the heliotropic behaviour in *A. viridis*, and we confirmed this at multiple stages: 1. Heliotropism in *A. viridis* aligns with the presence of the endosymbiont **(Fig. 1H-J)**. 2. Heliotropic responses are triggered by light wavelengths that are associated with photosynthetic activity **(Fig. 1A & B)**. 3. Blocking photosynthesis in symbiotic *A. viridis* has impaired heliotropism **(Fig 4D & E)**. 4. By measuring the rate of O_2_ production, we demonstrated a direct correlation between the rate of photosynthesis and the magnitude of heliotropic responses **(Fig 4F & G)**.

Though our results indicate a direct correlation between heliotropism and the rate of photosynthesis in *A. viridis*, that could be interpreted as a way to enhance photosynthetic efficiency. However, the morphological aspects of *A. viridis* under heliotropism do not concur with this hypothesis. We observed that by aligning its tentacles parallel to the incident light direction, *A. viridis* reduced the surface area of the tissue exposed to solar irradiation by ∼ 40% **(Fig 4H & I)**. This raises the question of whether heliotropism in *A. viridis* is an adaptive mechanism to promote light absorption or conversely to minimise excess light uptake.

In plants, the leaves display two kinds of heliotropism movements: diaheliotropism and paraheliotropism. Under diaheliotropism, the leaf blade orients perpendicularly to the incident sun’s light direction to increase light capture **(Fig 4J)**. In contrast, paraheliotropism (light avoiding) acts to orient the leaf blade parallel to the incident light to reduce the absorption of solar irradiances **(Fig 4J)** [52-59]. Similarly, pointing tentacles parallel to the incident sunlight in *A. viridis* is likely a regulatory mechanism that modulates the influx of solar irradiance reaching the endosymbionts **(Fig 4K)**. The paraheliotropic movement in plant leaves is principally determined by photo-oxidative stress due to photoinhibition (photosynthesis saturation) and water loss via transpiration due to rising surface temperature [52-59]. In plants and other photosynthetic organisms like algae, excess light accelerates the production of reactive oxygen species (ROS), resulting in photodamage. Therefore, to avoid net photoinhibition, certain plants utilise paraheliotropic movement to reduce light absorption to prevent photodamage [54, 55]. Like plants, the motile green algae exhibit negative phototaxis under increased solar irradiation to avoid photodamage [60]. Unlike free-living algae, endosymbiotic algae are circumscribed to the cnidarian host cell microenvironment. Therefore, photosynthetically produced O_2_ by endosymbionts can diffuse into the surrounding host cells and potentially increase reactive ROS production by the host [61, 62]. Indeed, photosynthetically produced O_2_ has been shown to be a controlling factor in locomotion/phototaxis in symbiotic sea anemones like *Anthopleura elegantissima* and *Aiptasia* [27, 41]; in these species, hyperoxia inhibited positive phototaxis. Previous studies in *A. viridis* revealed the presence of high levels of antioxidant enzymes in both host and symbiont cells, and the maintenance of antioxidant enzyme and catalase activities is directly proportional to endosymbiont O_2_ production capacity [63, 64]. However, high solar irradiance can overcome the ability of the host antioxidant scavenging system to sufficiently eliminate undesirable ROS formation and prolonged oxidative stress can trigger the early stages of symbiosis breakdown, resulting in bleaching [65-67].

The evidence from measured O_2_ production in *A. viridis* suggests that under bright sunlight, both hyperoxia and heliotropism occur simultaneously **(Fig 4G, 3A & B)**. Additionally, the measurements of the electron transfer rate (ETR: the quantum yield of photosynthesis photon flux density) of isolated *Symbiodinium* from *A. viridis* showed significant photoinhibition at higher irradiance values **(Fig. 4K)**, suggesting an excess light exposure can lead to photoinhibition in endosymbionts. The level of oxidative damage depends on the interplay of ROS production, antioxidant capacity, and cellular repair processes. Therefore, additional host strategies and their interplay with symbionts at the cellular level can directly affect the ability of the host to cope with oxidative stress and thereby indirectly determine the threshold of bleaching in symbiotic cnidaria [68, 69]. Hence, to maintain the stable symbiotic state, the endosymbionts and host anemones have likely evolved mutually beneficial behavioural mechanisms such as phototaxis [18, 28]. Similarly, *A. viridis* pointing their tentacles towards the sun’s position could be a regulatory system that modulates the influx of solar irradiance reaching the endosymbionts and filters out the excess solar irradiance that can damage the endosymbiont’s photosynthesis, which is similar to the paraheliotropism strategy witnessed in plants **(Fig 4J)**. Heliotropism in symbiotic *A. viridis* may have provided an advantage for colonising a wide range of light-exposed habitats, from shallow rock pools with the highest mid-day solar irradiance to deep waters and caves where light is scarce [32].

Although we widely recognise the mutually beneficial relationship between cnidarian hosts and their photosynthetic endosymbionts, the mechanisms of interactions and the impact of changes in the endosymbiont on the host are not fully understood. Our study suggests that there may be spontaneous signalling between the host and endosymbiont; in this case, the endosymbiont influences the anemone’s behaviour. While our data suggests potential factors like ROS contributing to heliotropism, we are still uncertain about how the endosymbionts and hosts communicate, which is fundamental to understanding the endosymbiotic relationship in cnidarians. Irrespective of the functional relevance of heliotropism in *A. viridis* either to promote light absorption or conversely to minimise excess light uptake, we revealed how photosymbiotic *A. viridis* displays similar behaviour as plants under similar environmental pressures.

## Materials and Methods

### The field experiment

The field experiment was conducted to examine the influence of sunlight on *A. viridis* behaviour. This study was conducted at Wembury, Plymouth, UK (50°19’00.5”N 4°05’00.6”W). We examined multiple rock pools and collected images. For heliotropism measurement, we placed a scale adjacent to *A. viridis* as a reference for the horizontal surface, and correspondingly, we also recorded the sun’s position. The angle of tentacle orientation was measured by superimposing a protractor over the collected images.

#### Anemonia viridis

*A. viridis* were collected from the vicinity of Plymouth, UK shores and transferred to an outdoor tank system. The outdoor tank system is supplied with a continuous flow of natural seawater at a flow rate = ∼6 litres/minute, filtered to 100 μm, salinity = 35 ± 1 ppt, temperature = 12-14 °C, and natural diurnal light cycle under sunlight. To obtain aposymbiotic (bleached) *A. viridis*, individuals were kept in constant darkness until they completely lost *Symbiodinium*. The absence of *Symbiodinium* in tentacles was validated using fluorescence microscopy. The bleached *A. viridis* were cultured in similar conditions as symbiotic *A. viridis*; the only difference is that the tanks were kept in the dark. However, bleached *A. viridis* were moved out of dark conditions to clean the tanks once a week. Both symbiotic and bleached *A. viridis* were fed 3-4 times weekly with freshly hatched *Artemia*. Over 60 symbiotic and 25 bleached *A. viridis* specimens were used in the current study. In our aquarium system, symbiotic and bleached *A. viridis* show similar levels of asexual reproduction, suggesting they are in good health. Additionally, both bleached and symbiotic *A. viridis* reacted similarly to *Artemia* feeding and mechanical stimuli like touch and vibration, confirming that the motility and sensitivity were not impaired in bleached anemones. Further, when we exposed bleached *A. viridis* to light like in symbiotic *A. viridis*, they showed similar immediate reactions; however, bleached *A. viridis* didn’t show a heliotropism-like response.

### Behavioural experiments

#### In-vitro heliotropism

*A. viridis* maintained under natural conditions were transferred to an aquarium filled with seawater at 12 -14 °C up to 8.5 cm depth. Before the experiment, *A. viridis* were left for 40 minutes in the dark to acclimatise to the new conditions. After 40 minutes, *A. viridis* were provided with cool white light (240 μmol m^-2^ s^-1^). To simulate the directional changes in sunlight, the light source was moved 90° to the adjacent side every 10 minutes in a clockwise direction. The experiment finished when the light returned to the initial position. During the whole experiment, a camera recorded the *A. viridis* behaviour.

#### Heliotropic response under different light wavelengths

We used a minimum of four symbiotic and four bleached *A. viridis* for the analysis. The *A. viridis* were transferred to the aquarium and left for 40 minutes to acclimate to the new conditions in the dark. After the acclimatisation period, *A. viridis* were exposed to cool white light for 7 minutes (240-μmol m^-2^ s^-1^). To monitor responses to light intensity, six symbiotic and six bleached *A. viridis* were exposed to 9 minutes-long exposure to two different cool white light intensities (14 and 240 μmol m^-2^ s^-1^) with 40 minutes-long acclimatisation in darkness and 40 minutes-long dark exposure in between the two light treatments. To monitor responses to different light wavelengths, six symbiotic and six bleached *A. viridis* specimens were exposed to violet (380-435nm), blue (435-490nm) or cyan/green light (490-520nm), provided by Fluval Marine Nano LED and Kessil lighting (Evolution Aqua Ltd). Each exposure lasted 9 minutes, followed by 10 minutes of a dark interval between subsequent exposures, allowing tentacles to relax to a random horizontal arrangement. Throughout our in-vitro behavioural experiments, we tested one individual at a time and collected the data. All behaviour experiments were conducted at the same time of the day. However, we tested multiple individuals in a single tank in the initial trial experiments while standardising the experiment setup.

#### Identifying opsins and cryptochromes in *A. viridis*

Total RNA was isolated from tentacles of six symbiotic and six bleached *A. viridis* specimens using TRI reagent® (Sigma) according to the manufacturer’s instructions. The concentration and quality of RNA were determined using a Nanodrop spectrometer (Thermo Scientific). cDNA was synthesised using the SuperScript® III Reverse Transcriptase (ThermoFisher) following the manufacturer’s protocol, and the concentrations were measured using a Nanodrop spectrometer. BLAST searches were performed on a representative transcriptome of *A. viridis* (GenBank under accessions SRS886541)[70] containing 1,164,058 contigs, using published cnidarian photoreceptor sequences [71]. Three opsin and three cryptochrome genes were used for the qPCR study. In the following list the gene name is first, followed by the NCBI GenBank Accession number (Opsin1, FK729339.1 forward 5’-ATACGCCCTCACCTCTCTGTAT-3’, reverse 5’-GGATGGCTATGCGGTAAAATGG-3’; Opsin2 FK733353.1 forward 5’-GCTGGGAATCGTCTCTCTCTAC-3’, reverse 5’-AGGAATATGCAAGGGGTGACAC-3’; Opsin3 FK759560.1: forward 5’-TCTCTTCGAAATTCAGGCATT-3’, reverse 5’-ACTTGTCCGTCATTTCTGTGG-3’; AnthoCRYI FK725676.1 forward 5’-GTTGTCAAAGTATCCTGTGCGC-3’, reverse 5’-GAGGACCCAACTTGCGGATAAT-3’; AnthoCRYII, FK735825.1: forward 5’-ACATGTGGTAGTCTGTGGGTTG-3’, reverse 5’-AGCTGTAACTCCTCAAACACCT-3’; AnthozoanCRY, FK725691.1: forward 5’-GATGGCTAACTCGCTGCATTG-3’, reverse 5’-CAACAGGCATCACACAAGAACC-3’; GAPDH FK723542.1 forward 5’-GAAGGTCATCATCTCAGCTC-3’, reverse 5’-GTGTAGCTGTATAGGCATGG-3’; 18S ribosomal FK727053.1 forward 5’-GTGACGGAGAATTAGGGTTC-3’, reverse 5’-CCTCCAATGGATCCTCGTTA-3’; ACTB FK727002.1 forward 5’-CATGTACGTAGCTATCCAGGC-3’, reverse 5’-CTGTGGTGGTGAAGGAGTAAC-3’) Primers were validated for the right product size and melt curves prior to qPCR. The qPCRs were run on Corbett Rotor-Gene RG-6000 Real-Time PCR Analyzer using a SensiFAST™ SYBR® No-ROX Kit (Meridian©). Gene encoding 18S was used as a housekeeping gene; prior to these, we compared the 18S, GAPDH and ACTB housekeeping genes and confirmed that 18S was the most stable expression gene.

#### DCMU treatment

To measure the effect of DCMU (3-(3,4-dichlorophenyl)-1,1-dimethylurea) on *Symbiodinium* photosynthesis, *Symbiodinium* from tentacles of four individuals were extracted. Five tentacles were slit open from each *A. viridis* with a scalpel to scrape the *Symbiodinium* from the *A. viridis* tissue and vortexed to obtain a homogenous *Symbiodinium* suspension. The *Symbiodinium* was then divided equally between tubes, each containing approximately 2–3 × 10^5^ cells/ml. Control and DCMU-treated samples were incubated in filtered seawater or filtered seawater with 10 or 100 μgl^-1^ DCMU. After 1 hour in the dark, Fv/Fm for each sample was measured using pulse amplitude modulated (PAM) chlorophyll fluorometry (Heinz Walz GmbH). In parallel, the Fv/Fm was measured on symbiotic tentacles that were collected from *A. viridis* exposed to different DCMU treatments, a minimum of four *A. viridis* were used in this experiment. Firstly, Fv/Fm was measured prior to the DCMU exposure, after one hour in 10 or 100 μgl^-1^ DCMU. Lastly, we measured how the effect of DCMU concentrations changes with time. Extracted *Symbiodinium* cells were incubated in four different DCMU concentrations: 0, 0.1, 1 and 10 μgl^-1^. Fv/Fm of all the samples was then measured using PAM after 1 and 24 hours.

#### DCMU treatment effect on *A. viridis* behaviour

Four symbiotic *A. viridis* were transferred to the aquarium of the size 14.5 × 14.5 × 14.5 cm filled with seawater of 12 -14 °C up to 8.5 cm high. LED light was placed above the aquarium, whilst the camera was placed on the side of the aquarium. *A. viridis* were first left for 10 minutes in the dark to acclimatise to the new conditions and then exposed for 5 minutes to cool white light (240 μmol m^-2^ s^-1^). After the data were collected for all four *A. viridis*, each of them was transferred into an individual beaker filled with seawater and DCMU (concentration: 100μgl^-1^) and exposed to ambient sunlight. The same step was then repeated after 2 and 24 hours of DCMU exposure.

### O_2_ Measurements

Fibre-optic O_2_ microsensors (PyroScience, Germany) (tip size of 50-70μm) were used to measure O_2_ production. The microsensor was connected to a FireSting®-O_2_ meter. The O_2_ sensor readings were linearly calibrated from measurements in air-saturated seawater and in anoxic seawater (produced by adding sodium sulfite to seawater at experimental temperature and salinity). O_2_ measurements were collected directly from the tissue surface of *A. viridis* tentacles. During each experiment, measurements were collected from at least ten tentacles per animal during 5 minutes of light exposure, and each tentacle was measured for approximately 30 seconds with a measurement sample interval of 1/sec. Micromanipulators and a stereo microscope were used to position the microsensor on the *A. viridis* tentacles.

### Data analysis

#### Behavioural studies

To analyse data collected on the behavioural response of *A. viridis* during the first phototactic, colour, and DCMU experiments, we used the program GIMP 2.10.18. Using the program, a template consisting of 10 sections (I-X) that were horizontally stacked on top of each other was created. The template was scaled according to each *A. viridis* size, with only the longest tentacles pointing straight up, reaching the top of the highest section (section X). The position of each tentacle was established at the section where the tip of the tentacle reached (only long tentacles growing nearest to the oral disc were included in the analysis). The number of tentacles was calculated for each of the sections at a given point, and the percentage of tentacles in each section at the given point was calculated. The differences in the response of symbiotic *A. viridis* towards different light colours and the difference in the response of symbiotic and bleached *A. viridis* to different light wavelengths were established by comparing the percentage of tentacles reaching the top two sections IX and X. *A. viridis* response to the white light throughout the course of 7 minutes and the effect of DCMU and its length of exposure on *A. viridis* behaviour was tested by comparing the percentage of tentacles reaching top 3 sections VIII, IX, X to cover the maxim number of tentacles. As for the intensity experiment, we compared the percentage of tentacles reaching Sections I-III, IV-VII, and VIII-X. When the light was positioned on the side, we calculated the percentage of the tentacles pointing in the direction of the light source at the given point.

All data from behavioural studies were analysed in R. To establish whether we have to carry out a parametric test or non-parametric equivalent of the test, we checked the assumption of equal variances/homogeneity of variance using the Levene’s test and whether the variables followed a normal distribution using Shapiro-Wilk test. Where two or more dependent samples were compared, we used either a paired t-test or One-way repeated Measures ANOVA (in case of heterogeneous data or not normally distributed data Friedman test). The Welch Two Sample t-test or Wilcoxon rank-sum test was run when comparing bleached and symbiotic *A. viridis*.

#### PAM analysis

Fv/Fm measured in filtered seawater was used as the highest possible photosynthetic performance for each *A. viridis*, and based on its value, we determined the performance in percentage for all the *Symbiodinium* samples extracted from the same *A. viridis* in different DCMU treatments. To analyse the effect of DCMU concentration on photosynthesis and how this effect changes over time, a Paired sample t-test was performed for normally distributed and homogeneous data, and the Asymptotic Wilcoxon signed-rank test for not normally distributed and heterogeneous data.

#### O_2_ analysis

The O_2_ measurements collected from all four individuals under specific light conditions (Dark, violet, blue and cyan) were averaged. A Two-tailed test was performed to assess the significance between dark and other light conditions, including violet, blue and cyan.

### Measuring surface Area

We used the ImageJ 1.52a (National Institute of Health, USA, http://imageJ.nih.Gov/ij) [72] tool to digitally measure the surface area of *A. viridis* tissue exposed towards the light through a threshold-based pixel count measurement to calculate the area of interest. This image analysis method was used in studies to digitally analyse regions of interest, such as leaf [73-75] or tissue areas [76, 77]. All images were taken from above the animal at a similar distance. Images were edited when necessary to remove shadows and darken light areas. After setting the scale (Analyse> Set Scale), the image was converted into 8-bit grayscale images (Image > Type > 8-bit). Next, the threshold was adjusted (Image > Adjust > Threshold) and was computed to measure the area (Analyse> Measure). The data was transferred to Excel for statistical analysis.

### *Symbiodinium* rapid light curves

Freshly isolated *Symbiodinium* from tentacles were dark acclimated for 30 min. The electron transfer rate was measured with a Water PAM (PAM WinControl 3) at a range of increasing light intensities (Heinz Walz GmbH). After the dark acclimation, measurements were collected in nine illumination steps: 29, 102, 153, 235, 346, 528, 786, 1111 and 1531 μmol m^-2^ s^-1^. Each illumination step lasted one min and was followed by a measurement of the effective quantum yield of PSII (UPSII = (F0m - Ft)/F0m), from which the PamWin software calculates the photosynthetic ETR as ETR =ΦII (quantum yield of photochemical energy conversion in PS II) x 0.84 × 0.5 x PAR [78]. The light curves are plots of ETRs versus actinic irradiance.

## Supporting information

Movie S1: Anemones react to the changing light conditions.

Movie S2: Anemone behavioural responses differ with the intensity of light.

Movie S3: A. viridis can track the light's movement when the light position changes every 10 minutes.

Movie S4: A. viridis display altered responses to different wavelengths of light.

Movie S5: Aposymbiotic anemones did not show altered responses to different wavelengths of light.

Movie S6: Aposymbiotic anemones did not show any ability to bend towards the light and track the movement of the light.

## Supplementary information

**Movie S1:** Anemones react to the changing light conditions. Temporary blocking of sunlight results in rapid loss of heliotropic behaviour, and similarly, upon restoring the sunlight, the tentacles point towards the direction of the sun.

**Movie S2:** Anemone behavioural responses differ with the intensity of light. The anemone exhibits a strong response under bright light conditions. Low: low-intensity light; High: high-intensity light.

**Movie S3:** *A. viridis* can track the light’s movement when the light position changes every 10 minutes. The tentacle’s orientation adjusts rapidly to changes in the light position.

**Movie S4:** *A. viridis* displays altered responses to different wavelengths of light. Blue and purple light showed a significantly higher effect on anemone heliotropic behaviour by pointing a large proportion of tentacles straight up compared to cyan light treatments.

**Movie S5:** Aposymbiotic anemones did not show altered responses to different wavelengths of light.

**Movie S6:** Aposymbiotic anemones did not show any ability to bend towards the light and track the movement of the light.

## Acknowledgements

We thank Kevin Atkins for his help with setting up the sea anemone facility. We thank Alix Harvey for their support in maintaining the animal facility.

## Author contributions

VM, EL and CB were involved in conceptualising the study; EL, VM and CB designed the experiments; CS and VM performed field experiments; VM and MK executed O_2_ experiments; EL performed all other experiments and analyses; EL and VM wrote the original draft; All authors were involved in reviewing and editing the text; Funding acquisition and project supervision: VM.

## References

1. Morey, K.J., et al., Chapter Twenty-Five - Developing a Synthetic Signal Transduction System in Plants, in Methods in Enzymology, C. Voigt, Editor. 2011, Academic Press. p. 581–602.

2. Gualtieri, P., Morphology of photoreceptor systems in microalgae. Micron, 2001. 32(4): p. 411–426.

3. Foster, K.W. and R.D. Smyth, Light Antennas in phototactic algae. Microbiol Rev, 1980. 44(4): p. 572–630.

4. Ueki, N., et al., Eyespot-dependent determination of the phototactic sign in Chlamydomonas reinhardtii. Proc Natl Acad Sci U S A, 2016. 113(19): p. 5299–304.

5. Kathare, P.K. and E. Huq, Signals | Light Signaling in Plants, in Encyclopedia of Biological Chemistry III (Third Edition), J. Jez, Editor. 2021, Elsevier: Oxford. p. 78–89.

6. Liscum, E., et al., Phototropism: growing towards an understanding of plant movement. The Plant cell, 2014. 26(1): p. 38–55.

7. Kami, C., et al., Light-regulated plant growth and development. Curr Top Dev Biol, 2010. 91: p. 29–66.

8. Corrochano, L.M. and V. Garre, Photobiology in the Zygomycota: Multiple photoreceptor genes for complex responses to light. Fungal Genetics and Biology, 2010. 47(11): p. 893–899.

9. Galland, P. and E.D. Lipson, Blue-light reception in Phycomyces phototropism: evidence for two photosystems operating in low- and high-intensity ranges. Proceedings of the National Academy of Sciences, 1987. 84(1): p. 104–108.

10. Loeb, J. and H. Wasteneys, On the Identity of Heliotropism in Animals and Plants. Proc Natl Acad Sci U S A, 1915. 1(1): p. 44–7.

11. Meltzer, S.J., The Dynamics of Living Matter. Science, 1906. 24(605): p. 145–7.

12. Thompson, D.A.W., Forced Movements, Tropisms, and Animal Conduct. Nature, 1919. 103(2583): p. 163–164.

13. Atamian, H.S., et al., Circadian regulation of sunflower heliotropism, floral orientation, and pollinator visits. Science, 2016. 353(6299): p. 587–90.

14. Zhang, S., et al., Flower heliotropism of Anemone rivularis (Ranunculaceae) in the Himalayas: effects on floral temperature and reproductive fitness. Plant Ecology, 2010. 209(2): p. 301–312.

15. Ehleringer, J. and I. Forseth, Solar tracking by plants. Science, 1980. 210(4474): p. 1094–8.

16. Sailaja, M.V. and V.S. Rama Das, Leaf solar tracking response exhibits diurnal constancy in photosystem II efficiency. Environmental and Experimental Botany, 1996. 36(4): p. 431–438.

17. Habermann, G., et al., Leaf paraheliotropism in Styrax camporum confers increased light use efficiency and advantageous photosynthetic responses rather than photoprotection. Environmental and Experimental Botany, 2011. 71(1): p. 10–17.

18. Foo, S.A., et al., Photo-movement in the sea anemone Aiptasia influenced by light quality and symbiotic association. Coral Reefs, 2020. 39(1): p. 47–54.

19. Shick, J.M. and W.I. Brown, Zooxanthellae-Produced O2 Promotes Sea-Anemone Expansion and Eliminates Oxygen Debt under Environmental Hypoxia .1. Journal of Experimental Zoology, 1977. 201(1): p. 149–155.

20. Pearse, V.B., Modification of Sea-Anemone Behavior by Symbiotic Zooxanthellae - Phototaxis. Biological Bulletin, 1974. 147(3): p. 630–640.

21. Sawyer, S.J., H.B. Dowse, and J.M. Shick, Neurophysiological Correlates of the Behavioral Response to Light in the Sea Anemone Anthopleura elegantissima. Biol Bull, 1994. 186(2): p. 195–201.

22. Fournier, A. The story of symbiosis with zooxanthellae, or how they enable their host to thrive in a nutrient poor environment.

23. Klaus, J.S., et al., Environmental controls on corallite morphology in the reef coral Montastraea annularis. Bulletin of Marine Science, 2007. 80(1): p. 233–260.

24. Kuhlmann, D.H.H., Composition and Ecology of Deep-Water Coral Associations. Helgolander Meeresuntersuchungen, 1983. 36(2): p. 183–204.

25. Anthony, K.R.N. and O. Hoegh-Guldberg, Variation in coral photosynthesis, respiration and growth characteristics in contrasting light microhabitats: an analogue to plants in forest gaps and understoreys? Functional Ecology, 2003. 17(2): p. 246–259.

26. Kishimoto, M., et al., Negative phototaxis in the photosymbiotic sea anemone Aiptasia as a potential strategy to protect symbionts from photodamage. Scientific Reports, 2023. 13(1): p. 17857.

27. Strumpen, N.F., et al., High light quantity suppresses locomotion in symbiotic Aiptasia. Symbiosis, 2022.

28. Shick, J.M., et al., Oxygen Uptake in Sea Anemones: Effects of Expansion, Contraction, and Exposure to Air and the Limitations of Diffusion. Physiological Zoology, 1979. 52(1): p. 50–62.

29. Pardy, R.L., Aspects of Light in the Biology of Green Hydra, in Coelenterate Ecology and Behavior, G.O. Mackie, Editor. 1976, Springer US: Boston, MA. p. 401–407.

30. Wilson, E.B., The Heliotropism of Hydra. The American Naturalist, 1891. 25(293): p. 413–433

31. Kim, S. and J.T. Robinson, Phototaxis is a state-dependent behavioral sequence in <em>Hydra vulgaris</em>. bioRxiv, 2023: p. 2023.05.12.540432.

32. Wiedenmann, J., C. Röcker, and W. Funke, The morphs of Anemonia aff. sulcata (Cnidaria, Anthozoa) in particular consideration of the ectodermal pigments, in Verhandlungen der Gesellschaft für Ökologie, J. Pfadenhauer, Editor. 1999, Spektrum Akademischer Verlag. p. 497–503.

33. Porro, B., et al., The many faced symbiotic snakelocks anemone (Anemonia viridis, Anthozoa): host and symbiont genetic differentiation among colour morphs. Heredity (Edinb), 2020. 124(2): p. 351–366.

34. Banerjee, R. and A. Batschauer, Plant blue-light receptors. Planta, 2005. 220(3): p. 498–502.

35. Christie, J.M. and W.R. Briggs, Blue light sensing in higher plants. J Biol Chem, 2001. 276(15): p. 11457–60.

36. Allemand, D. and P. Furla, How does an animal behave like a plant? Physiological and molecular adaptations of zooxanthellae and their hosts to symbiosis. Comptes Rendus Biologies, 2018. 341(5): p. 276–280.

37. Bricaud, A., et al., Natural variability of phytoplanktonic absorption in oceanic waters: Influence of the size structure of algal populations. Journal of Geophysical Research-Oceans, 2004. 109(C11).

38. Kinzie, R.A., P.L. Jokiel, and R. York, Effects of light of altered spectral composition on coral zooxanthellae associations and on zooxanthellae in vitro. Marine Biology, 1984. 78(3): p. 239–248.

39. Levy, O., Z. Dubinsky, and Y. Achituv, Photobehavior of stony corals: responses to light spectra and intensity. Journal of Experimental Biology, 2003. 206(22): p. 4041–4049.

40. Moya, A.l., et al., Study of calcification during a daily cycle of the coral Stylophora pistillata: implications for ‘light-enhanced calcification’. Journal of Experimental Biology, 2006. 209(17): p. 3413–3419.

41. Fredericks, C.A., Oxygen as a limiting factor in phototaxis and in intraclonal spacing of the sea anemoneAnthopleura elegantissima. Marine Biology, 1976. 10 Vol. 38;(Iss. 1).

42. Leach, W.B. and A.M. Reitzel, Decoupling behavioral and transcriptional responses to color in an eyeless cnidarian. BMC Genomics, 2020. 21(1): p. 361.

43. Plachetzki, D.C., C.R. Fong, and T.H. Oakley, Cnidocyte discharge is regulated by light and opsin-mediated phototransduction. BMC Biology, 2012. 10(1): p. 17.

44. Artigas, G.Q., et al., A gonad-expressed opsin mediates light-induced spawning in the jellyfish Clytia. Elife, 2018. 7.

45. North, W.J., Sensitivity to light in the sea anemone Metridium senile (L). II. Studies of reaction time variability and the effects of changes in light intensity and temperature. J Gen Physiol, 1957. 40(5): p. 715–33.

46. Wang, J.T., et al., Diverse responses of Symbiodinium types to menthol and DCMU treatment. PeerJ, 2017. 5: p. e3843.

47. Fransolet, D., S. Roberty, and J.-C. Plumier, Impairment of symbiont photosynthesis increases host cell proliferation in the epidermis of the sea anemone Aiptasia pallida. Marine Biology, 2014. 161(8): p. 1735–1743.

48. Watanabe, T., I. Yuyama, and S. Yasumura, Toxicological effects of biocides on symbiotic and aposymbiotic juveniles of the hermatypic coral Acropora tenuis. Journal of Experimental Marine Biology and Ecology, 2006. 339: p. 177–188.

49. Parrin, A. and N. Blackstone, THE ROLE OF LIGHT AND PHOTO-OXIDATIVE STRESS IN CORAL BLEACHING. The FASEB Journal, 2017. 31(S1): p. 889.11–889.11.

50. Vogelmann, T.C. and L.O. Björn, Response to directional light by leaves of a sun-tracking lupine (Lupinus succulentus). Physiologia Plantarum, 1983. 59(4): p. 533–538.

51. Wainwright, C.M., SUN-TRACKING AND RELATED LEAF MOVEMENTS IN A DESERT LUPINE (LUPINUS ARIZONICUS). American Journal of Botany, 1977. 64(8): p. 1032–1041.

52. Chávez, R.O., et al., Detecting Leaf Pulvinar Movements on NDVI Time Series of Desert Trees: A New Approach for Water Stress Detection. PLOS ONE, 2014. 9(9): p. e106613.

53. Zhu, C.G., et al., Heliotropic leaf movement of Sophora alopecuroides L.: An efficient strategy to optimise photochemical performance. Photosynthetica, 2015. 53(2): p. 231–240.

54. Takahashi, S. and M.R. Badger, Photoprotection in plants: a new light on photosystem II damage. Trends in Plant Science, 2011. 16(1): p. 53–60.

55. Arena, C., L. Vitale, and A.V. De Santo. Paraheliotropism in Robinia pseudoacacia Plants: An Efficient Means to Cope with Photoinhibition. in Photosynthesis. Energy from the Sun. 2008. Dordrecht: Springer Netherlands.

56. Levizou, E. and A. Kyparissis, A novel pattern of leaf movement: the case of Capparis spinosa L. Tree Physiology, 2016. 36(9): p. 1117–1126.

57. Jayawardena, D.M., et al., Elevated carbon dioxide plus chronic warming causes dramatic increases in leaf angle in tomato, which correlates with reduced plant growth. Plant, Cell & Environment, 2019. 42(4): p. 1247–1256.

58. Koller, D., Light-driven leaf movements*. Plant, Cell & Environment, 1990. 13(7): p. 615–632.

59. Pastenes, C., et al., Paraheliotropism can protect water-stressed bean (Phaseolus vulgaris L.) plants against photoinhibition. Journal of Plant Physiology, 2004. 161(12): p. 1315–1323.

60. Wakabayashi, K.-i., et al., Reduction-oxidation poise regulates the sign of phototaxis in <i>Chlamydomonas reinhardtii</i>. Proceedings of the National Academy of Sciences, 2011. 108(27): p. 11280–11284.

61. Dykens, J.A., et al., Oxygen Radical Production in the Sea Anemone Anthopleura Elegantissima and its Endosymbiotic Algae. Journal of Experimental Biology, 1992. 168(1): p. 219–241.

62. Mydlarz, L.D. and R.S. Jacobs, An inducible release of reactive oxygen radicals in four species of gorgonian corals. Marine and Freshwater Behaviour and Physiology, 2006. 39(2): p. 143–152.

63. Richier, S., et al., Characterization of superoxide dismutases in anoxia- and hyperoxia-tolerant symbiotic cnidarians. Biochimica et Biophysica Acta (BBA) - General Subjects, 2003. 1621(1): p. 84–91.

64. Richier, S., et al., Symbiosis-induced adaptation to oxidative stress. Journal of Experimental Biology, 2005. 208(2): p. 277–285.

65. Richier, S., et al., Oxidative stress and apoptotic events during thermal stress in the symbiotic sea anemone, Anemonia viridis. The FEBS Journal, 2006. 273(18): p. 4186–4198.

66. Lesser, M.P., Oxidative stress causes coral bleaching during exposure to elevated temperatures. Coral Reefs, 1997. 16(3): p. 187–192.

67. Downs, C.A., et al., Oxidative stress and seasonal coral bleaching. Free Radical Biology and Medicine, 2002. 33(4): p. 533–543.

68. Rädecker, N., et al., Heat stress destabilizes symbiotic nutrient cycling in corals. Proceedings of the National Academy of Sciences, 2021. 118(5): p. e2022653118.

69. Weis, V.M., Cellular mechanisms of Cnidarian bleaching: stress causes the collapse of symbiosis. Journal of Experimental Biology, 2008. 211(19): p. 3059–3066.

70. Macrander, J., M. Broe, and M. Daly, Tissue-Specific Venom Composition and Differential Gene Expression in Sea Anemones. Genome Biol Evol, 2016. 8(8): p. 2358–75.

71. Gornik, S.G., et al., Photoreceptor Diversification Accompanies the Evolution of Anthozoa. Molecular Biology and Evolution, 2020. 38(5): p. 1744–1760.

72. Schroeder, A.B., et al., The ImageJ ecosystem: Open-source software for image visualization, processing, and analysis. Protein Sci, 2021. 30(1): p. 234–249.

73. Orsini, F., et al., A comparative study of salt tolerance parameters in 11 wild relatives of Arabidopsis thaliana. J Exp Bot, 2010. 61(13): p. 3787–98.

74. Warman, L., A.T. Moles, and W. Edwards, Not so simple after all: searching for ecological advantages of compound leaves. Oikos, 2011. 120(6): p. 813–821.

75. Juneau, K.J. and C.S. Tarasoff, Leaf Area and Water Content Changes after Permanent and Temporary Storage. PLOS ONE, 2012. 7(8): p. e42604.

76. Modepalli, V., et al., Marsupial tammar wallaby delivers milk bioactives to altricial pouch young to support lung development. Mech Dev, 2016. 142: p. 22–29.

77. Gómez Martín, C. and G. Martínez Grau, Use of ImageJ as an image processing method for the assessment of post-surgical bruises. Skin Research and Technology, 2021. 27(5): p. 655–667.

78. Baker, N.R., Chlorophyll fluorescence: a probe of photosynthesis in vivo. Annu Rev Plant Biol, 2008. 59: p. 89–113.

